# Assessing the biogeography of marine giant viruses in four oceanic transects

**DOI:** 10.1101/2023.01.30.526306

**Authors:** Anh D. Ha, Mohammad Moniruzzaman, Frank O. Aylward

**Author notes:** **Correspondence:** Frank O. Aylward.

## Abstract

Viruses of the phylum *Nucleocytoviricota* are ubiquitous in ocean waters and play important roles in shaping the dynamics of marine ecosystems. In this study, we leveraged the bioGEOTRACES metagenomic dataset collected across the Atlantic and Pacific Oceans to investigate the biogeography of these viruses in marine environments. We identified 330 viral genomes, including 212 in the order *Imitervirales* and 54 in the order *Algavirales*. We found that most viruses appeared to be prevalent in shallow waters (<150 meters), and that viruses of the *Mesomimiviridae* (*Imitervirales*) and *Prasinoviridae* (*Algavirales*) are by far the most abundant and diverse groups in our survey. Five mesomimiviruses and one prasinovirus are particularly widespread in oligotrophic waters; annotation of these genomes revealed common stress response systems, photosynthesis-associated genes, and oxidative stress modulation that may be key to their broad distribution in the pelagic ocean. We identified a latitudinal pattern in viral diversity in one cruise that traversed the North and South Atlantic Ocean, with viral diversity peaking at high latitudes of the northern hemisphere. Community analyses revealed three distinct *Nucleocytoviricota* communities across latitudes, categorized by latitudinal distance towards the equator. Our results contribute to the understanding of the biogeography of these viruses in marine systems.

## Introduction

Large DNA viruses of the phylum *Nucleocytoviricota*, also known as “giant viruses”, are a diverse group of eukaryotic viruses with particle sizes typically larger than 0.2 μm in diameter and genome sizes reaching up to 2.5 Mbp [1–4]. Known members of the phylum are partitioned into six orders, namely *Algalvirales, Imitevirales, Pimascovirales, Pandoravirales, Asfuvirales*, and *Chitovirales*, and up to 32 potential families [5]. These viruses have an ancient origin and have underwent frequent gene exchange with their hosts [6, 7], and as a result their genomes often encode numerous genes involved in cellular processes such as glycolysis, the TCA cycle, amino acid metabolism, translation, light sensing, and cytoskeletal dynamics [8–15]. Giant viruses are known to infect a broad spectrum of eukaryotic hosts; while members of the *Imitervirales, Algavirales*, and *Pandoravirales* infect a wide range of algae and various heterotrophic protists, members of the *Asfuvirales, Chitovirales*, and *Pimascovirales* infect a wide range of protist and metazoan hosts [2, 16–19]. The diverse functional repertoires harbored in these viruses’ genomes are thought to render them able to manipulate the physiology and subvert the immune responses of their hosts during infection [9, 20, 21]. Giant viruses therefore play important roles in various ecological processes across the globe.

Although giant viruses are ubiquitous in the biosphere, they appear to be particularly abundant and diverse in marine environments. Early studies focusing on amplification and sequencing of the viral Family B DNA polymerase from seawater found that algal viruses within the *Nucleocytoviricota* were widespread in a variety of marine environments [22, 23], an observation which was later confirmed through analysis of community metagenomic data [24, 25]. A recent comparative metagenomic study found that giant viruses are ubiquitous in the ocean, vary markedly across depth, and are prevalent in >0.22 um size fractions [26]. Field studies have estimated that the abundance of giant viruses can reach up to 10^4^ −10^6^ viruses per milliliter of seawater, with higher abundances typically recovered during algal blooms [27–32]. Giant viruses have been reported to infect many prevalent marine eukaryotic lineages, including chlorophytes, haptophytes, and choanoflagellates [15, 18, 33, 34], and they are therefore an important factor shaping marine ecological dynamics. Moreover, several studies have shown that giant viruses associated with algal blooms play key roles in carbon export to deeper waters [35–37], indicating they are critical components of global carbon cycles. Despite the ecological importance of giant viruses, our understanding of their diversity lags behind that of smaller viruses owing to the widespread use of filtration steps in viral diversity surveys, which often exclude larger viruses [38]. There is therefore a strong need for further studies to examine the biogeography and ecological dynamics of these large viruses in the ocean.

Here, we aimed to undertake a genome-based global survey of giant virus assemblages across the oceans and compare the diversity of these viruses in different geographic regions by leveraging the large number of metagenome-assembled genomes (MAGs) of these viruses that have been generated [9, 39–41]. We focused on the metagenomic data generated from samples collected in four transects in the Atlantic and Pacific oceans as part of the the GEOTRACES project [42], which provide clear, well-defined geographic and depth profiles. These metagenomes targeted the >0.2 um size fraction and are therefore suitable for examining the diversity of large viruses. Our work broadens our understanding of the biogeography of giant viruses in the ocean and reveals novel diversity patterns that will be important for understanding their role in marine environments.

## Results and Discussion

### Imitervirales and Algavirales orders are the most abundant and diverse among viruses recovered in bioGEOTRACES metagenomic data set

We examined 480 metagenomes collected and sequenced as part of the bioGEOTRACES component of the GEOTRACES project [42]. These samples were derived from four major cruises from different regions of the South Pacific and Atlantic Oceans (Fig. 1) in 2010-2011. This dataset targeted the >0.2-μm size fraction of microbial communities and was sampled along well-defined transects at various depths, making it suitable for assessing the geographic and depth distribution of marine giant viruses. In total, we identified 330 giant virus genomes with metagenomic reads mapping. To investigate the taxonomic distribution of the detected *Nucleocytoviricota* viruses, we constructed a phylogenetic tree of the viruses and 1,188 *Nucleocytoviricota* reference genomes (Fig. S1; see Methods). Out of the 330 genomes recovered, we were able to place the genomes within the orders *Imitervirales* [n=214], *Algavirales* [n=54], *Pimascovirales* [n=16], *Pandoravirales* [n=4], and *Asfuvirales* [n=1]. On the family level, the most well-represented groups were the *Mesomimiviridae* [n=146] and *Prasinoviridae* [n=42] (Fig. 2). We also identified 41 *Mirusviricota* viruses; although technically not members of the *Nucleocytoviricota*, these large DNA viruses are prevalent in the ocean and share many genomic features with giant viruses [41]. Of the 330 genomes identified, only 8 were derived from cultivated viruses, and 322 genomes are metagenome-assembled genomes (MAGs). Approximately half (159) of the 330 genomes are larger than 300 kbp (Table S1), underscoring the prevalence of viruses with large genomes throughout the ocean.

**Figure 1.**
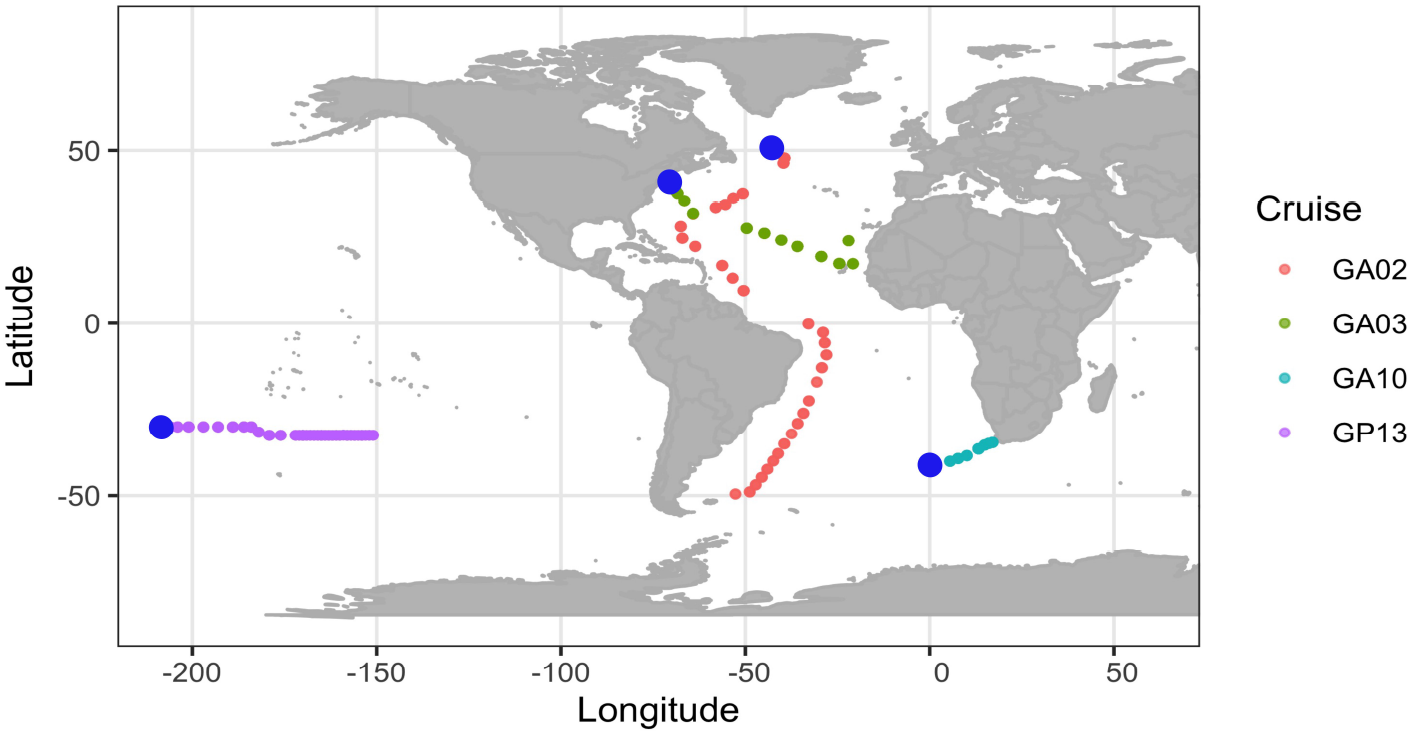
Global map of sampling locations, colored by transect. Blue dots indicate the start of cruise tracks.

**Figure 2.**
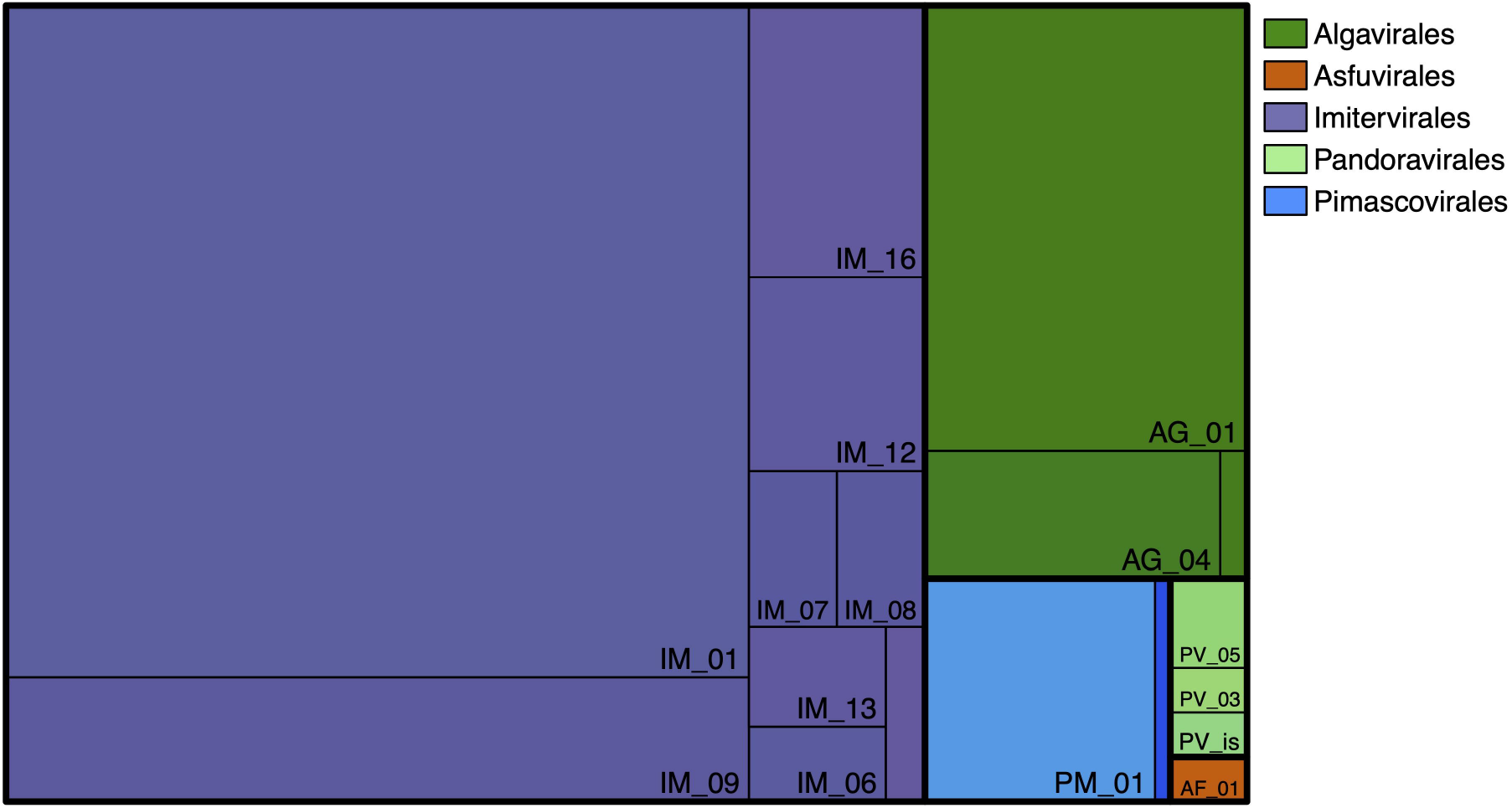
Summary of the taxonomy of detected giant viruses. The area of each rectangle is proportional to the number of identified viral genomes in the respective taxon.

Of the 330 genomes with reads mapping, 14 viruses, including 8 *Algavirales* and 5 *Imitervirales* were recovered in all four bioGEOTRACES transects. Meanwhile, 182 viruses were found in only one transect, among which 116 genomes were found exclusively in transect GA02, most of which were *Imitervirales* (Fig. 3A, Table S1). In term of total giant virus richness, the number of different genomes found in transect GA02, which traces along the Americas-Atlantic Ocean coastline (248 genomes total) by far exceeded that in the other three transects, especially compared to the pelagic transect GP13 where less than one-third of that number (71 genomes) were detected. This is likely a consequence of the much broader range of latitudes and biogeochemical regimes sampled by the GA02 transect compared to the others. Across the transects, 65 viruses were present in all three depth layers of the water columns (<80m, 80-150m, and >150m), most of which were *Imitervirales* and *Algavirales* (Fig. 3B). We found 103 viruses that solely appeared in the surface water of 80 meters up, while there were only 5 genomes unique to the deep water of below 150 meters.

**Figure 3.**
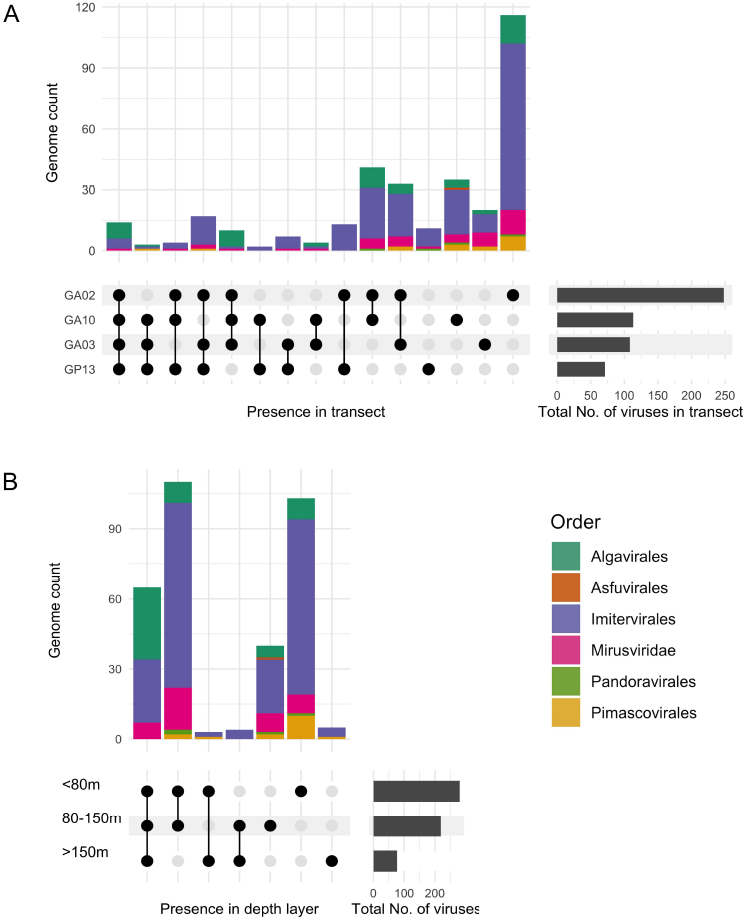
Unique genomes and genomes shared between the transects (A) and water depth layers (B). Horizontal bars (right) indicate the total number of genomes found in each transect; black dots indicate the presence in one or multiple transects; the corresponding vertical bars indicate the number of genomes with the presence described by the dots.

Giant virus communities were mostly dominated by members of the *Imitervirales* and *Algavirales* orders, regardless of the transect location or depth of sampling (Fig. 4). On average, *Imitervirales* and *Algavirales* accounted for 56.4% and 32.6% of the total number of giant virus occurrences across all sampling locations, respectively. This result is consistent with previous observations that viruses of these two orders were the most abundant and widespread giant viruses in the Pacific and Atlantic Ocean [25, 26, 28]. Viruses within the *Imitervirales* were particularly widespread in communities across all depths sampled in the pelagic GP13 transect, with a mean contribution of 88.8%, and the majority (9) of the 11 viruses found exclusively in GP13 were *Imitervirales* (Fig. 2A; Fig. 4C). This pattern of *Imitervirales* dominance in pelagic waters was also observed in transect GA03 (Fig. 4B, inner samples), underscoring the prevalence of this group in oligotrophic gyres. In general, the spatial distribution of viruses in the ocean is shaped largely by the geographic distribution of their hosts [43], and the broad distribution of viruses within the *Imitervirales* is therefore likely a signature of their collective broad host range. Indeed, members of the *Imitervirales* order are known to infect an exceptionally broad phylogenetic range of hosts [19], including marine haptophytes in the genera *Phaeocystis* and *Chrysochromulina*, which were found in high abundance in the open waters of the central Pacific Ocean [44, 45], as well as other widespread hosts such as the green algae, Choanoflagellates, and amoeboid protists [15, 46–49]. Aside from these established hosts, recent work using co-occurrence analyses have also identified a wide range of other potential eukaryotic hosts for viruses within the *Imitervirales* [12, 50, 51], suggesting that the hosts of viruses in this order is far broader than currently known. Lastly, given that many members of the *Imitervirales* gain entry to host cells through phagocytosis, it is likely that individual viral populations may infect a range of different host lineages. If this is the case, the broad representation of the *Imitervirales* in pelagic surface waters may represent the ability of viruses in this order to infect a range of mixotrophic and heterotrophic lineages.

**Figure 4.**
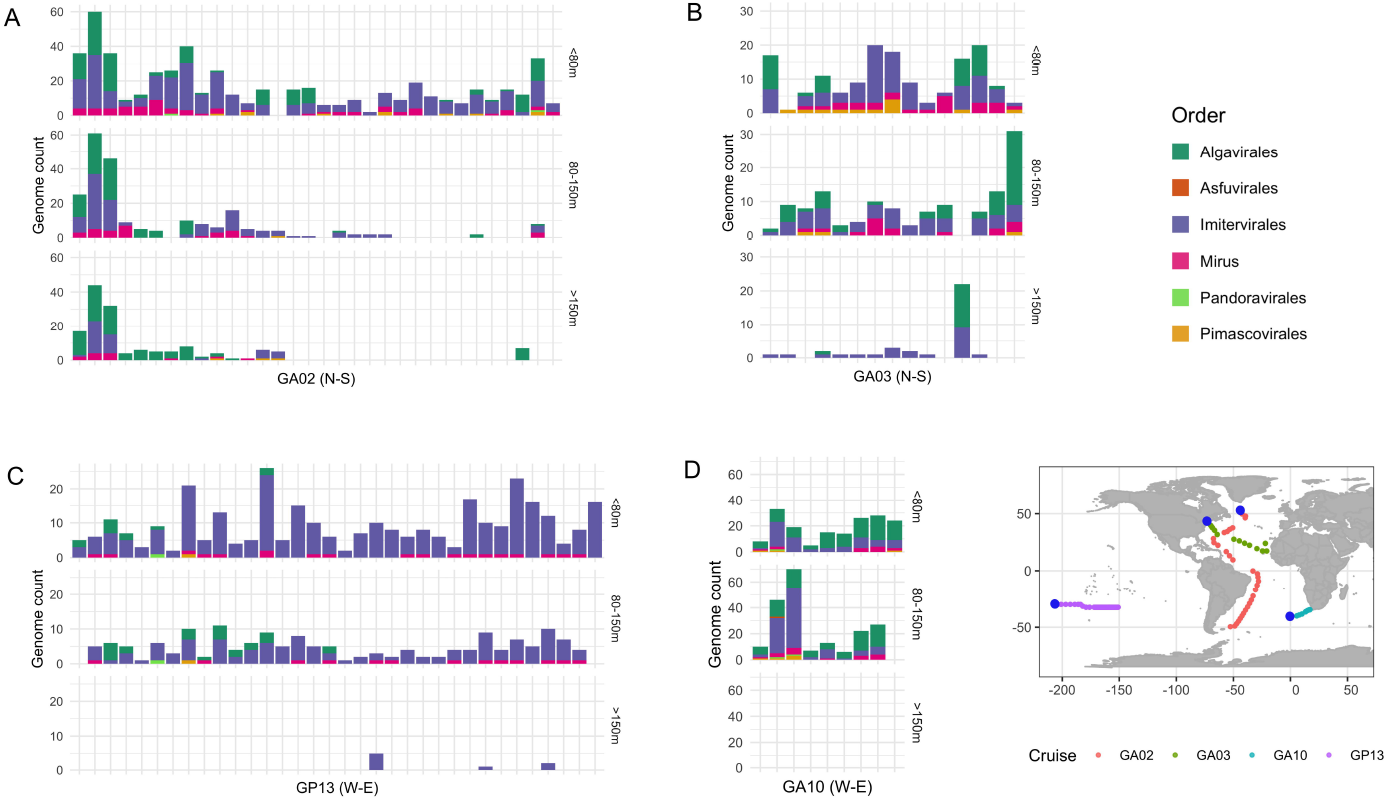
Distribution of giant viruses in each transect. Each column represents a sampling location. The y-axis shows the number of different viral genomes that were recovered at a given location, separated into three depth ranges (2-80m, 80-150m, and 150-5,500m). Locations are arranged in increasing distance from left to right on the x-axis, based on their distance from the starting location and follow the indicated orientation (N to S for GA02 and GA03, W to E for GA10 and GP13).

The majority of viruses that are most widespread in the Atlantic Ocean in our survey (i.e., found in all three Atlantic transects GA02, GA03, and GA10, but not in the Pacific transect GP13) were *Algavirales* viruses (Fig. 3A). Viruses within the *Algavirales* order were especially abundant in the North Atlantic Ocean (northern samples in transect GA02) and in the coastal waters (eastern samples in transect GA10) where we observed a high abundance throughout the water column (Fig. 4A, 4D; Fig. 5C). Viruses of this order were also present in high abundance in surface waters (<50m) and euphotic waters at 50-150m deep on both two coastal sides of the transect GA03, while showing a sharp decreasing trend towards the pelagic waters of the North Atlantic and the Pacific Ocean (inners of transect GA03 and transect GP13, respectively). Once again, these changes in the abundance of viruses within the *Algavirales* likely reflect the distribution of their hosts. Members of the family *Prasinoviridae* are the most prevalent family we identified within the *Algavirales*, and members of this group are known to infect members of the prasinophyte genera *Ostreococcus, Bathycoccus*, and *Micromonas* [18], which have been found to be highly abundant in coastal systems. It was estimated that prasinophytes may account for 50-90% of total picoeukaryotic cells in coastal waters, while they only made up a much lower fraction (<20%) of those in pelagic waters [52]. Although not as abundant as their counterparts in coastal populations, there are several prasinophytes found widespread in oligotrophic waters of the open ocean, for instance *O. lucimarinus* and *Micromonas spp*., allowing the broad presence of viruses infecting these hosts, which have been detected and documented [53, 54].

**Figure 5.**
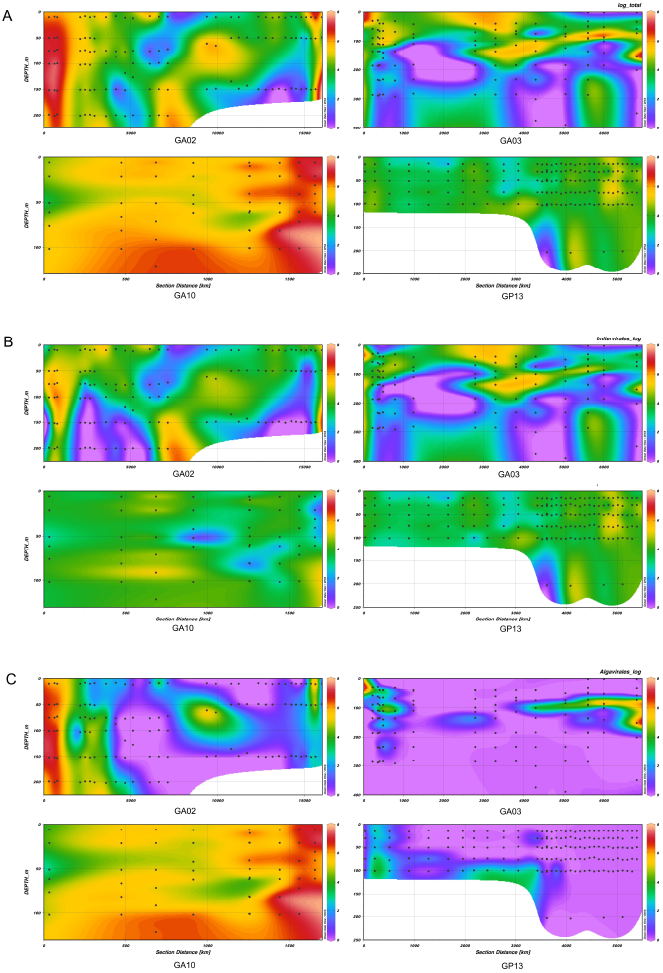
Distribution of viruses throughout the water column along the transects. The viral abundance (calculated in log RPKM) of (A) total giant viruses present in the transect (B) viruses of the *Imitervirales* order (C) viruses of the *Algavirales* order only. Samples were ordered based on the distance along transects, beginning from the first sampling location of cruise tracks (0 km). Black dots denote the sampling location along the transect of each sample.

Mirusviruses, *Pimascovirales, Pandoravirales*, and *Asfuvirales* were present to a lesser degree in the four transects, with average contributions of 9.1%, 1.5%, 0.3%, and 0.03% of giant virus occurrence, respectively. Mirusviruses were prevalent across all four transects (Fig. 4), consistent with findings of a previous investigation using metagenomic read recruitments from Tara Oceans datasets [41]. These viruses appeared to be more abundant in coastal waters, which was maintained throughout the water column (transect GA02 and east of transect GA10), while in pelagic waters their abundance was more limited to sunlit waters at <100m (inners of transect GA03 and transect GP13, respectively) (Fig. S2A). This pattern is in agreement with the prediction that Mirusviruses infect a broad planktonic host range that includes many phototrophs [41]. All of the *Pimascovirales* genomes recovered in our survey were MAGs derived from marine metagenomes. Although currently little is known about the natural hosts of this viral group in the oceans, their prevalence across the four transects suggests that they infect widespread host taxa and play important roles in marine systems. Interestingly, we found that the recently-delineated family-level clade PM_01 was the most prevalent lineage of the *Pimascovirales* in the ocean, but no members of this group have been cultivated and their host range remains unknown. A recent study found a member of this lineage was prevalent in surface waters of Station ALOHA, consistent with the view that they are present in oligotrophic surface waters [55]. The only *Asfuvirales* virus found in our survey, GVMAG-M-3300027833-19 was recovered in the GA10 transect, which is located off the Atlantic coast of South Africa. This viral genome was also recently recovered in a TARA metagenomic sample in the same region [17] and its transcriptomic activities were detected in the waters of the central California Current upwelling system in the North Pacific Ocean [12], suggesting that the virus may be widely distributed beyond the sampling scope of the bioGEOTRACES cruises. *Pandoravirales* viruses were present in all four transects in both coastal and pelagic waters, although at relatively low number of occurrences and abundance. Across all locations included in our survey, their distribution was strictly limited to shallower waters (<150 meters) (Fig. S2C). The *Pandoravirales* includes well-studied coccolithoviruses that are prevalent in *Emiliania huxleyi* blooms, suggesting that at least some members of this order will have highly variable abundance depending on host availability. An important caveat of our study is that we surveyed only metagenomes derived from >0.2 um size fractions; it is likely that some members of the *Pimascovirales* and *Asfuvirales*, which have on average smaller genome and virion sizes than members of the *Imitervirales*, may be more widespread in smaller size fractions, and are therefore more prevalent than indicated by our results here.

Giant viruses were apparently more diverse and abundant in surface waters (Fig. 3B, Fig. 4). Indeed, of all occurrences of giant viruses throughout the water column at all locations, 92.1% were located in waters at 150 meters or shallower. On average across all four transects, the viral richness at two depth ranges (<80m and 80-150m) were approximately 2.7 and 2.3 times higher than that at deep waters (>= 150m), respectively (Kruskal–Wallis and Dunn’s test, P<0.001) (Fig. S3). In transects GA03 and GP13, the decline in total giant virus abundance with depth was particularly sharp (Fig. 5A). In deeper pelagic water (>200m), the giant communities appeared to be limited to just a few members of the *Imitervirales* (inners of transect GA03 and transect GP13), while in coastal waters at deeper than 200m, members of the *Algavirales* are also present with relatively high abundance, together with *Imitervirales* viruses (Fig. 4, Fig. 5). Apart from viruses of the *Algavirales* and *Imitervirales* orders, our read mapping approach did not recover any giant virus MAGs of any other orders in metagenomes sequenced from water sampled from depths >200m. This may be partially due to biases in the reference database, which includes more genomes from surface water samples, but it seems likely that it is at least partially driven by the large diversity of giant viruses in surface waters.

### Latitudinal pattern of giant virus diversity

We analyzed the giant virus communities along the transect GA02 in more detail to assess possible latitudinal gradients in giant virus diversity. The GA02 transect sampling sites follows the Americas-Atlantic Ocean coastline, spanning across a long range of latitudes from the parallel 50° North to 50° South and tracing a clear latitudinal gradient from the North Atlantic in the summer of 2010 to the south Atlantic in the austral summer of 2011. This sampling scheme may facilitate the detection of subtle latitudinal gradients that may be more difficult to resolve through comparison of samples collected across different ocean basins. In terms of community composition at each sampling location, the *Algavirales* assemblages seemingly dominated the giant virus communities in northern samples and decreased in abundance towards the south, replaced by the dominance of viruses of the order *Imitervirales* (Fig. S4).

We calculated taxonomic richness and Shannon’s H diversity index in each depth-integrated sampling location to investigate latitudinal variation (Fig. 6). To avoid biases in diversity measurements due to unequal sequencing depth, we rarefied all metagenomic samples to 10M reads prior to calculation. We detected a latitudinal pattern of diversity along the GA02 transect with average diversity increasing with higher latitudes in the Northern Hemisphere and plateaued towards the South. Total giant virus communities peaked, both in terms of richness and alpha diversity in the further north of the North Atlantic Ocean (i.e. above 40° North) and steeply declined around the middle latitudes (20-40° North). The trend of increasing diversity from the equatorial zone towards higher latitudes was mirrored in the *Imitervirales* communities, while varying marginally in the *Algavirales* communities for both viral genome richness and alpha diversity (Fig. 6). The clear peak in latitudinal diversity in the Northern Hemisphere is consistent with the trend of species richness observed for a large portion of the total of 65,000 marine species examined previously [56]. It is possible that stronger environmental instability, particularly the wide temperature variation in the northern hemisphere (excluding polar zones) [57] may explain the higher diversity compared to the south. A relatively similar northern spike of giant virus diversity has been reported from analyses of the Family B DNA Polymerase (PolB) genes in Tara Oceans datasets [26, 58], although the studies observed another increase in diversity near the southern middle latitudes. The discrepancy did not seem to result from the disparity in methodological approaches between our mapping strategy and the above two studies; we also performed calculation of diversity indices on the TARA Ocean datasets using our mapping method described herein, and observed a similar trend in giant virus diversity agreeing with in the two PolB studies in latitudinal locations of elevated diversity (Fig. S5). It is possible that the slightly differing results reflects the fact that the bioGEOTRACES and Tara Oceans samples were collected from different times with different cruise tracks.

**Figure 6.**
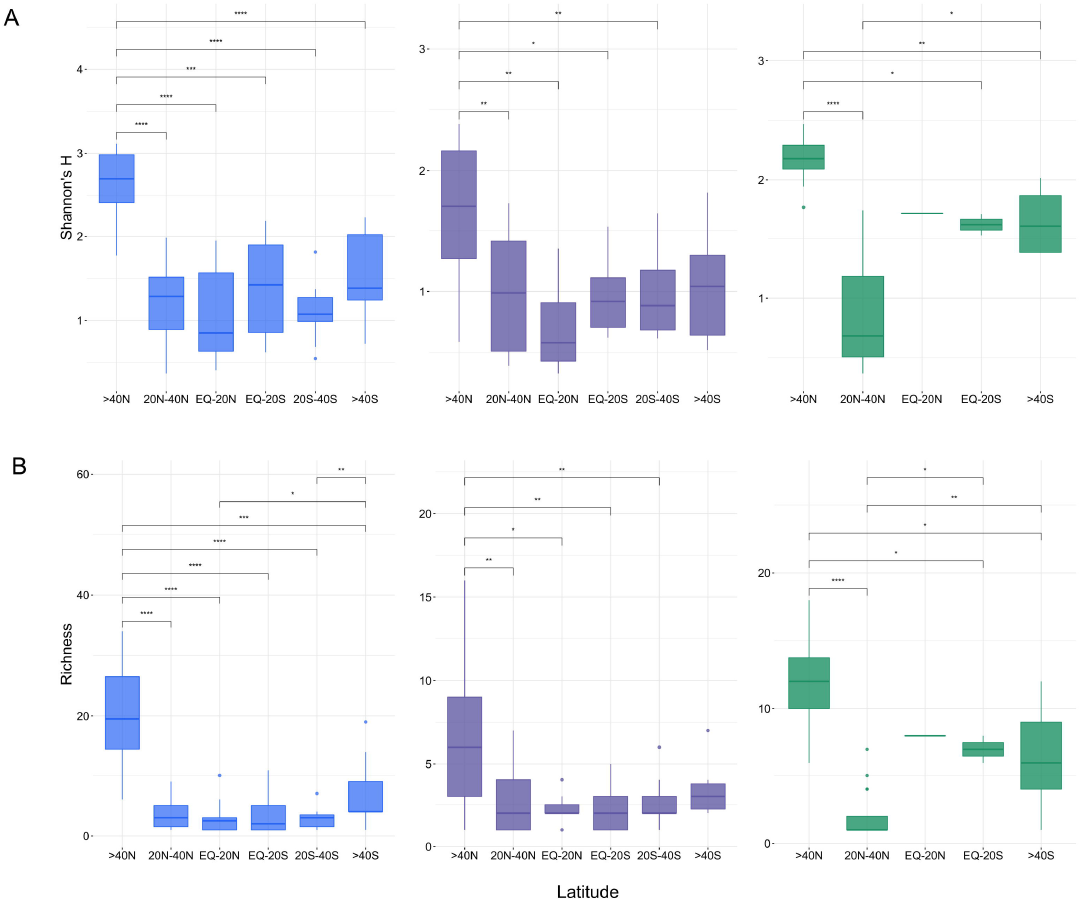
Latitudinal pattern of giant virus diversity across the transect GA02 showed in (A) Shannon’s H index (B) Genome richness. Stars showing significant difference between two latitudinal groups (Wilcox test, p-values < 0.05) (* < 0.05, ** < 0.01, *** < 0.001, **** < 0.0001) Panels left: Total virus community; center: Imitervirales communities; right: Algavirales communities. EQ, Equator.

The lower diversity, in terms of alpha diversity and richness of the giant virus communities near the equator in comparison with northern high latitudes did now follow conventional latitude diversity pattern, which posits that marine eukaryotic diversity generally increases toward the tropics [58, 59]. The difference could possibly be attributed to several potential causes. High temperature in the equatorial zone is potentially one underlying cause; decline in species diversity at higher temperatures has been observed in several marine taxa [60]. Previous work has widely reported that marine species tend to shift away from the tropics to higher latitudes due to climate warming [61–63]. A species distribution model has projected that marine species may acquire latitudinal peaks in total richness at approximately 40/30° absolute latitude and a loss of richness near the equator, highlighting the important role of temperature on these distribution shifts [64]. Indeed, the metagenomic samples located in the GA02 transect included in our survey were collected during the months from late March to June, which were anticipated to be the warmest months of the year in the equatorial Atlantic Ocean [65]. Given the strong influence of seasonality on marine microbial communities, it is likely that latitudinal gradients in diversity are ephemeral and will vary throughout the year.

A non-metric multidimensional scaling (NMDS) analysis indicated that *Nucleocytoviricota* communities were clustered according to their latitudinal distance to the equator (Fig. 7). The giant virus community composition significantly differed between three latitudinal sectors in both the GA02 transect exclusively and all transects collectively (Permanova p < 0.001). This clustering is fairly consistent with traditional Longhurst oceanographic biogeographical biomes of plankton ecology, which were designated based on the distribution of chlorophyll, angle of sunlight, and cloudiness [66]. This may further support the view that the geographic distribution of viruses in ocean waters is mostly affected by the distribution of their hosts. In terms of viral community richness, we found only 5 giant virus genomes (2% of the total number of giant viruses found in the GA02 transect) shared across all three latitudinal zones (Fig. S6), all of which belonged to the order *Algavirales*. The high latitude zone (>40° latitude) harbored 93 unique genomes, while the mid-latitude zone (20° to 40° latitude) had 68 and the low latitude zone (between 20° equatorial) had 24.

**Figure 7.**
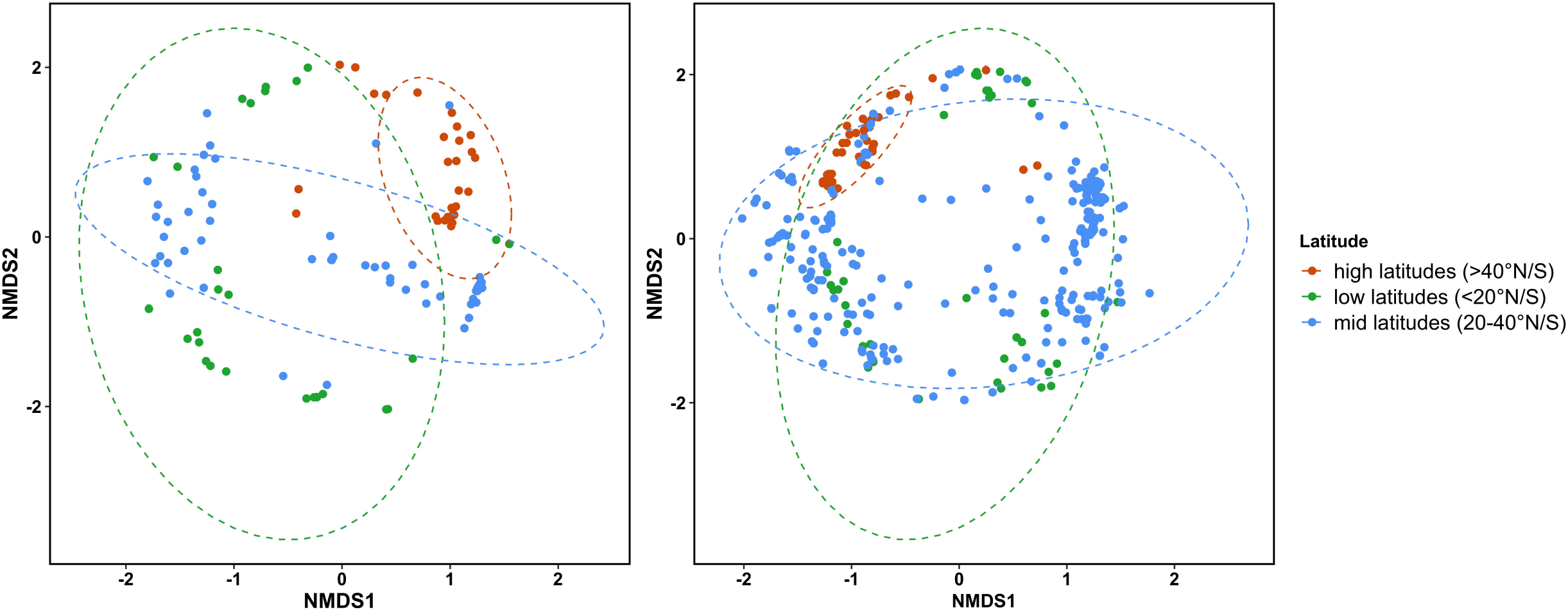
Community composition between latitudinal locations NMDS ordination based on Bray-Curtis distance matrices of viral communities collected in the (A) GA02 transect, stress = 0.3 (B) All four bioGEOTRACES, stress = 0.21. Latitudinal groups are color-coded by sample locations at higher than 40°N/S, from 20° to 40°N/S, and below 20°N/S (equatorial). Ellipses represent 95% confidence intervals. Viral communities are significantly different between groups (Permanova p <0.001).

### *Five* Mesomimiviruses *and one* prasinovirus *are particularly widespread in oligotrophic waters*

The vast majority of the most abundant and widespread viruses in our survey belong to the orders *Imitervirales* and *Algavirales*. We observed six MAGs within the *Imitervirales* (5 genomes) and the *Algavirales* (1 genome) that were particularly widespread in oligotrophic waters (Fig. 8A), all detected across different water depths in at least 19 distinct sampling locations. All the five *Imitervirales* could be classified into the recently-proposed *Mesomimiviridae* family (*Imitervirales* family 1) (Table S1). The *Mesomimiviridae* family is particularly widespread in marine systems and contains well-documented cultivated representatives that infect oceanic haptophytes, such as *Phaeocystis globosa* virus (PgV), *Chrysochromulina ericina* virus (CeV), and *Chrysochromulina parva* virus (CpV) [67–69]. Other members of the family have been found co-occurring and correlating with diatoms [50], suggesting that diatoms are potential hosts of this viral lineage. The only *Algavirales* virus belonged to the *Prasinoviridae* family (*Algavirales* family 1). The most broadly distributed genome, ERX556088.18.dc was recovered in 122 samples (more than 25% of the total number of samples analyzed overall) at 34 sampling locations. All the five Mesomimiviruses were extensively distributed in the Pacific transect GP13, while the prasinovirus, TARA_IOS_NCLDV_00011, was more widespread in the Atlantic Ocean (Fig. S7). All of these six genomes derived from marine environments, with genome sizes ranging from 108,412 bp to 483,524 bp and GC content varied from 26.2% to 34.6%.

**Figure 8.**
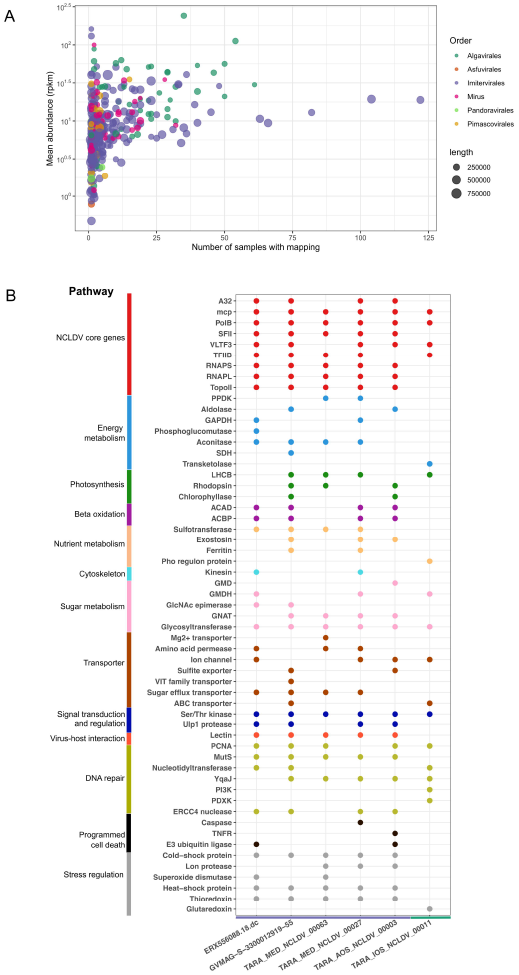
(A) General mapping statistics of viruses found in surface waters <150m. The y-axis shows the average abundance of a given individual genome (in RPKM), the x-axis shows the number of samples from which the virus was recovered. Dots are colored by the viral order and dot sizes represent the length of the genomes. (B) Genomic functional features of the six genomes that are widespread in oligotrophic waters. On the x axis, genomes are ranked from left to right in order of decreasing number of samples in which the viruses were detected; the horizontal colored bar shows the taxonomic order of the genome (purple: *Imitervirales*, green: *Algavirales*). The y axis denotes the functional annotation found in genomes; putative genes are color-coded by functional categories. Gene function abbreviations: PPDK, Pyruvate phosphate dikinase; GAPDH, Glyceraldehyde 3-P dehydrogenase; SDH, Succinate dehydrogenase; LHCB, Chlorophyll a/b binding protein; ACAD, Acyl-CoA dehydrogenase; ACBP, Acyl-CoA binding protein; GMD, GDP-mannose dehydrogenase; GMDH, GDP-mannose 4,6 dehydratase; GlcNAc epimerase, UDP-N-acetylglucosamine 2-epimerase; GNAT, Glucosamine-6-phosphate N-acetyltransferase; PCNA, Proliferating cell nuclear antigen; PI3K, Phosphatidylinositol 3-kinases; PDXK, PD-(D/E)XK nuclease superfamily; TNFR, Tumor necrosis factor receptor.

Annotation of these genomes showed complex genomic repertoires, which is a common characteristic of viruses of the *Mesomimiviridae* and the *Prasinoviridae*. Complete or near-complete set of 9 giant virus core genes, including major capsid protein (MCP), A32-like packaging ATPase (A32), superfamily II helicase (SFII), family B DNA Polymerase (PolB), virus late transcription factor 3 (VLTF3), large and small RNA polymerase subunits (RNAPL and RNAPS, respectively), TFIIB transcriptional factor (TFIIB), and Topoisomerase family II (TopoII) were found in all of the *Mesomimiviridae* genomes, indicating that these are high quality genome assemblies (Fig. 8B). These core genes are broadly represented in genomes of *Nucleocytoviricota* and have previously been used as phylogenetic markers for these viruses [5, 7, 70]. Both RNAP subunits were absent in the *Prasinoviridae* genome, consistent with the lack of DNA-dependent RNA polymerase that has been previously reported for prasinoviruses [11].

Other genes encoding essential viral functions were also consistently found in these genomes, including ribonucleotide reductase, thymidylate synthase, dUTPase (for nucleotide metabolism), Nudix-like hydrolase, mRNA capping enzyme (transcription and RNA processing), and glycosyltransferase (virion morphogenesis) (Table S2).

Genes involved in translation have been widely reported in the genomes of viruses within the order *Imitervirales* [71–73]. We found several translation-related genes, including aminoacyl-tRNA synthetases, or aaRS (asparaginyl-tRNA synthetase), translation initiation factors (IF4E, eIF3, IF1A), translation elongation factors (eF-TU) in all of the *Mesomimiviridae* genomes. The aaRS genes catalyzes the linkage between tRNAs and amino acids during translation and may act as a mechanism for circumventing nutrient starvation in the host cell, allowing the virus to maintain viral replication in different nutritional conditions [74].

Throughout all six genomes, we also identified numerous genes involved in diverse metabolic processes (Fig. 8B, Table S2), which may be involved in rewiring host metabolism and cellular physiology during infection to support their own viral production. We found genes involved in central carbon metabolism, including enzymes for glycolysis, the TCA cycle, and beta oxidation in all of the genomes. Numerous genes involved in nutrient acquisition and processing, light-driven energy generation, and diverse transporters were also present, consistent with previous findings [9, 39]. Rhodopsins could potentially alter the host’s sunlight-dependent energy transfer system [15], while chlorophyll a/b binding proteins might help maintain a stable light-harvesting capacity of host cells during infection [9]. The presence of genes involved in photosynthetic processes might be important for these viruses to infect a wide array of phototrophic or mixotrophic hosts in well-lit waters across the ocean. Genes encoding storage proteins and transporters, including ferritin-like proteins, amino acid permeases, transporters predicted to target sulfur, phosphorus, and iron are common in these genomes and may have a role in rewiring host’s nutrient acquisition strategies to enhance viral propagation. Such set of viral-encoded nutrient storage and transporters might be especially advantageous in marine environments, particularly in the oligotrophic waters of the South Pacific Ocean, where micronutrients such as iron are scarce [75] and the viruses need to employ their own transporters to boost nutrient acquisition. We also found homologs of genes involved in the regulation of cellular apoptosis, including caspase and tumor necrosis factor receptor. Manipulation of cell death is a common strategy employed by giant viruses to avert the impending cellular response to viral infection [76–78].

A broad array of stress response and repair genes found in all of the six genomes potentially equips the viruses with the ability to endure various external stresses common in oligotrophic waters, such as high temperatures, ultraviolet (UV) damage, and oxidative stress. We found genes involved in oxidative stress regulation, including thioredoxin, glutaredoxin, and superoxide dismutase (SOD) to be common among all genomes. Thioredoxin and SOD have been found expressed in several members of the *Imitervirales* [9, 79, 80] and were suggested to mitigate cellular oxidative stress by detoxifying harmful reactive oxygen species released by hosts during viral infection. SOD may also play an active role in reducing superoxide accumulation induced by UV exposures in direct sunlight, which may aid survival of viruses in the sunlit open waters [81]. It has been postulated that such viral-encoded redox genes allow the virus to infect a broad range of hosts [80]. Previous work has noted that giant viruses may carry genes that serve their own DNA repair to maintain high fidelity in genome replication [70, 82, 83]. We identified various DNA repair genes, including MutS mismatch repair and ultraviolet (UV) damage repair, such as ERCC4 nuclease [84] to be present in all of the mesomimivirus genomes. MutS homologs are widely present in genomes of mimivirus relatives [67, 85, 86] and are thought to associate with correcting mismatches to ensure the fidelity of viral genome replication. ERCC4-type repair nuclease might provide the viruses with crucial protection against DNA damages caused by UV irradiation. Although the prasinovirus’ genome lack homologs of MutS and ERCC4-type repair nuclease, it encodes numerous other putative DNA repair genes such as phosphatidylinositol 3-kinase and PD-(D/E)XK nuclease superfamily, which could also potentially aid in maintaining DNA integrity.

We also identified various genes predicted to encode enzymes for synthesizing glycans, which may be involved in the decoration of capsids with sugar moieties. These viral-encoded fibril structures are potentially useful for viruses to extend their host range and persist in the open waters. First, the oligosaccharides may enable the modification of virion surface to mimic the host’s normal food source, e.g. organic debris and bacteria [87, 88], promoting phagocytosis of virion particles. This strategy of infection, which takes advantage of the ‘generalized’ feeding habit that many marine protists rely on, may obviate the requirement of building receptors to a specific host and thus allow for a broader array of hosts. In addition, glycosylated fibrils could possibly act as a protective layer to shield the viruses from unfavorable environmental conditions, therefore increasing viral persistence. Furthermore, a study of Acanthamoeba polyphaga mimivirus has found that viral particles covered with self-produced sugars are able to adhere to different organisms through glycoside interactions, including bacteria, fungi, and arthropods [89], without infecting them. These organisms thus may help disperse the viruses over a wide area of waters, increasing their chance of contact with drifted host cells and expanding spatial distribution across the ocean. We also observed that all of these viruses carry lectin-domain containing proteins, which may act as key mediators of host-virus recognitions and interactions [90, 91]. Although the exact role of the protein in viruses is still unclear, it is possible that they might leverage lectin domains to modulate interactions with hosts and achieve a broader host range.

## Conclusion

In this study we conducted a metagenomic survey of giant viruses in the Atlantic and Pacific Oceans using the bioGEOTRACES datasets. We show that giant viruses of the orders *Imitervirales* and *Algavirales* are particularly widespread and abundant in epipelagic waters. Giant virus communities vary markedly by latitude, and in the GA02 transect in the Atlantic Ocean we detected a latitudinal pattern of diversity that peaks at high northern latitudes and plateaus towards the south. Lastly, we identified five genomes of the *Mesomimiviridae* family of the *Imitervirales* and one genome of the *Prasinoviridae* of the *Algavirales* that are particularly widespread in oligotrophic waters. Our comparative genomic analysis revealed that these genomes encoded diverse genes involved in central carbon metabolism, stress responses, and lectin-domain proteins potentially involved in host-virus interactions. We hypothesize that these genes may collectively expand the host range of these viruses, possibly explaining their particularly broad distribution. Overall, our study provides genomic insights into the distribution of giant viruses in the ocean and sheds light on the biogeography of these ecologically-important community members.

## Materials and Methods

### Nucleocytoviricota genome database compilation

We downloaded 1,382 *Nucleocytoviricota* genomes from the Giant Virus Database [5] and 696 viral MAGs assembled from 937 Tara Oceans metagenomes within the Global Ocean Eukaryotic Viral database [41]. All of these genomes were classified to the phylum *Nucleocytoviricota*, except those of the recently-discovered *Mirusviricota* lineage, which has a herpesvirus-like capsid and likely belongs to the realm Duplodnaviria. Although Mirusviruses represent a lineage distinct from the *Nucleocytoviricota*, we included them here because they represent a widespread lineage of marine large DNA viruses, and their genomes appear to be a chimera of different viral lineages, including the *Nucleocytoviricota*. To remove possible contamination from cellular sources, we screened all viral genomes using ViralRecall [92] and removed all contigs that had a score < 0 (indicating stronger signals from cellular sources). We also excluded genomes of less than 100 kbp total sequence, not encoding PolB gene, and/or containing less than 2 out of 4 of the marker genes SFII, TFIIB, VLTF3, and A32. To avoid the presence of identical or highly similar genomes, we dereplicated the genome set with dRep v3.2.2 [93] using an average nucleotide identity threshold of 95%. We arrived at a database containing 1,629 viral genomes (1,518 *Nucleoviricota* and 111 *Mirusviricota*) for metagenomic read mapping.

### Metagenome data set

We examined the metagenomic data from the >0.2-μm size fraction microbial communities of 480 samples collected by the international GEOTRACES program from May 2010 to December 2011. Accession numbers of the data are listed in Table S3. The samples were collected in four major cruise transects (GA02, GA03, GA10, and GP13) across the Atlantic and Pacific Oceans at 2-10 depths in each sampling location, ranging from 6m to 5601m. Sample processing was previously described in detail [42]. We calculated the geographical distance between sample locations in each transect based on recorded latitudes and longitudes using the function distHaversine from the R package geosphere.

### Reads processing and mapping

We downloaded and trimmed reads from each of the metagenome samples with Trim Galore v. 0.6.4 using parameters ‘–length 50 -e 0.1 -q 5 –stringency 1 --phred33”. We then mapped the trimmed reads onto the *Nucleocytoviricota* nucleotide sequences using coverM v0.6.1 (https://github.com/wwood/CoverM) in mode ‘genome’, with the parameter --min-read-percent-identity 0.95. We calculated relative abundance in reads mapped per kilobase of genome, per million mapped reads (RPKM). To avoid the false detection of viral genomes due to spurious read mapping, we only retained genomes with breadth coverage >20% (i.e. more than 20% of the genome length were covered by any read) in subsequent analyses. This cutoff is based on recent work which suggested that a genome coverage of at least 20% is appropriate to indicate the presence of that genome in a sample [94]. After this filtering, we obtained a set of 330 *Nucleocytoviricota* genomes for subsequent analysis.

### Phylogeny and clade delineation

To provide phylogenetic context for the giant virus genomes that we identified, we constructed a multilocus phylogenetic tree of the *Imitervirales* order using a set of 7 marker genes: family B DNA Polymerase (PolB), A32-like packaging ATPase (A32), Poxvirus late transcription factor 3 (VLTF3), superfamily II helicase (SFII), alpha RNA polymerase subunits (RNAPL), TFIIB transcriptional factor (TFIIB), and Topoisomerase family II (TopoII). The concatenated alignment of these 7 markers was generated with the ncldv_markersearch.py script (github.com/faylward/ncldv_markersearch) and then trimmed with TrimAl v. 1.4.rev22 [95] (parameter -gt 0.1). The tree was inferred from the alignment using IQ-TREE version 2.2.0.3 [96] with the best fitting model determined by the ModelFinder Plus option in IQ-TREE, according to the Bayesian Information Criterion (BIC). We used the same order-, family-, and genus-level nomenclature for the *Nucleocytoviricota* as previously described [5].

### Subsampling reads and calculating diversity

Comparison of diversity among samples, especially alpha diversity, may be erroneous due to differing library sizes [97]. To ensure equal library sizes across samples for diversity measurements, all samples were randomly subsampled without replacement to 10M reads using the reformat program provided in bbtools suite (Bushnell B. – sourceforge.net/projects/bbmap/). The subsampled reads were mapped against the viral genome set using coverM as described above. We then calculated community richness and Shannon’s diversity indices using the package ‘vegan’ (https://cran.r-project.org/web/packages/vegan/). Variation among community composition was analyzed with NMDS ordination based on Bray–Curtis dissimilarity using the function ‘metaMDS’, parameters k = 2, trymax = 100. Statistical analyses of difference in community composition were performed using a PERMANOVA test with the ‘adonis’ function, 9,999 permutations.

### Depth distribution mapping and interpolation

We performed interpolation of viral depth distribution using the program Ocean Data View v5.6.0 [98] in DIVA gridding mode, with parameters signal-to-noise = 25, automatic scale lengths for the X- and Y-axis, quality limit = 3 to exclude bad estimates.

### Protein annotations

We annotated proteins in six widespread giant virus genomes by comparing them to the EggNOG database 5.0.0 [99] and Pfam-A release 34 [100] hidden Markov models (HMMs) profile using HMMER v3.3.2 (parameter “-E 1e-3” for the EggNOG search and “–cut_nc” for the Pfam search) and retained only the best hits.

## Data availability

The data sets analyzed in this study are already publicly available and were accessed as described in the Materials and Methods section.

## Acknowledgements

We would like to thank the authors of Biller et al for providing access to the GEOTRACES metagenomes used in this study. We acknowledge the use of the Virginia Tech Advanced Research Computing Center for bioinformatic analyses performed in this study.

