## Supplementary Figures for "Assessing the biogeography of marine giant viruses in four oceanic transects"

**Figure S1. Phylogeny of all giant viruses used for metagenomic mapping.** The order-level classification of *Nucleocytoviricota* viruses is denoted by the branch colors and color strip. The genomes with mapping to transect metagenomic data are denoted on the outermost bar plot, with the height of the bar corresponding to genome size.

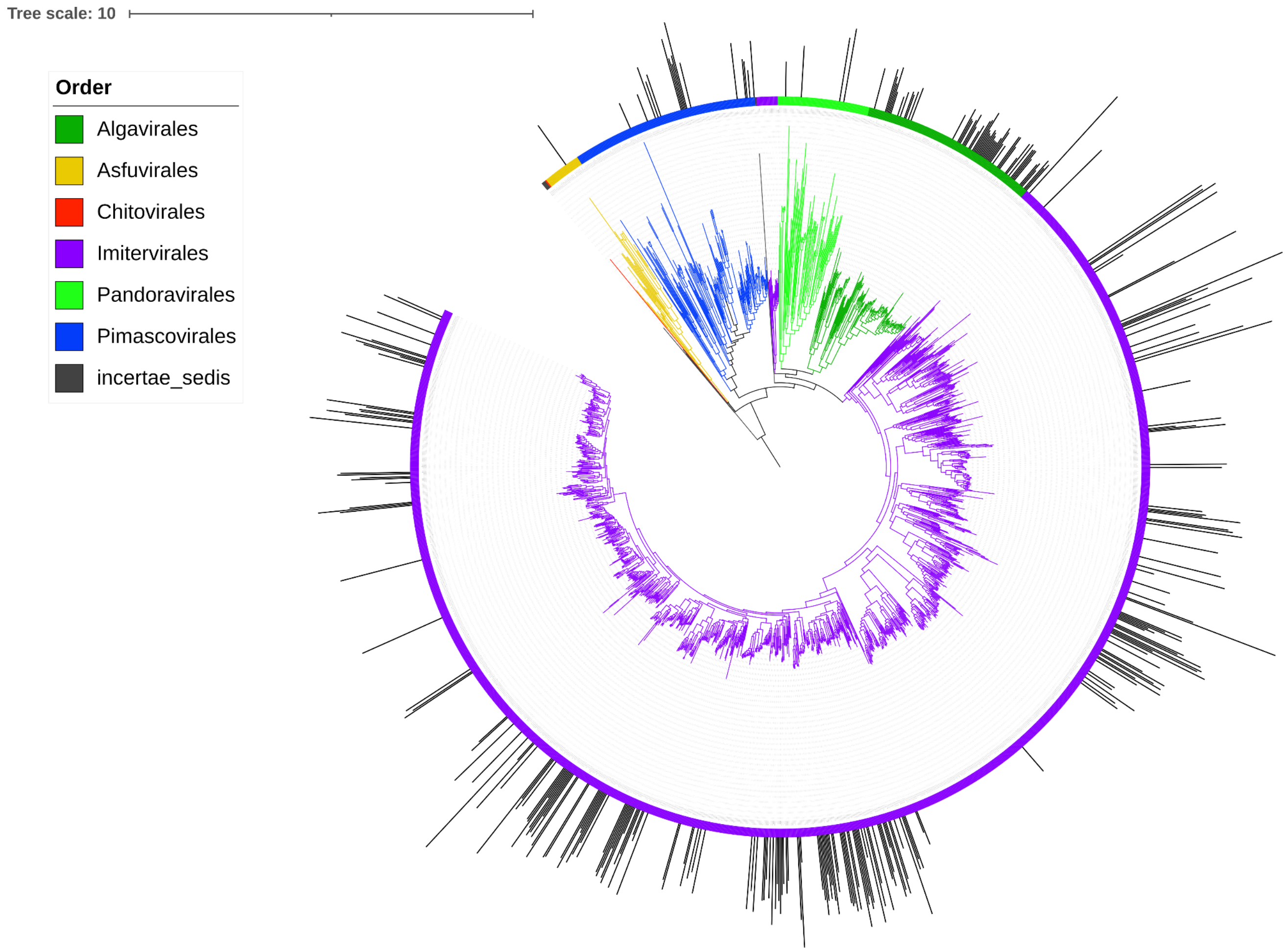

**Figure S2. Distribution of viruses of the *Mirusviricota* and the order *Pimascovirales* and *Pandoravirales* throughout the water column along the transects.** The viral abundance (calculated in log RPKM) of (A) *Mirusviricota* viruses (B) viruses of the *Pimascovirales* order (C) viruses of the *Pandoravirales* order. Samples were ordered based on the distance along transects, beginning from the first sampling location of cruise tracks (0 km). Black dots denote the sampling location along the transect of each sample.

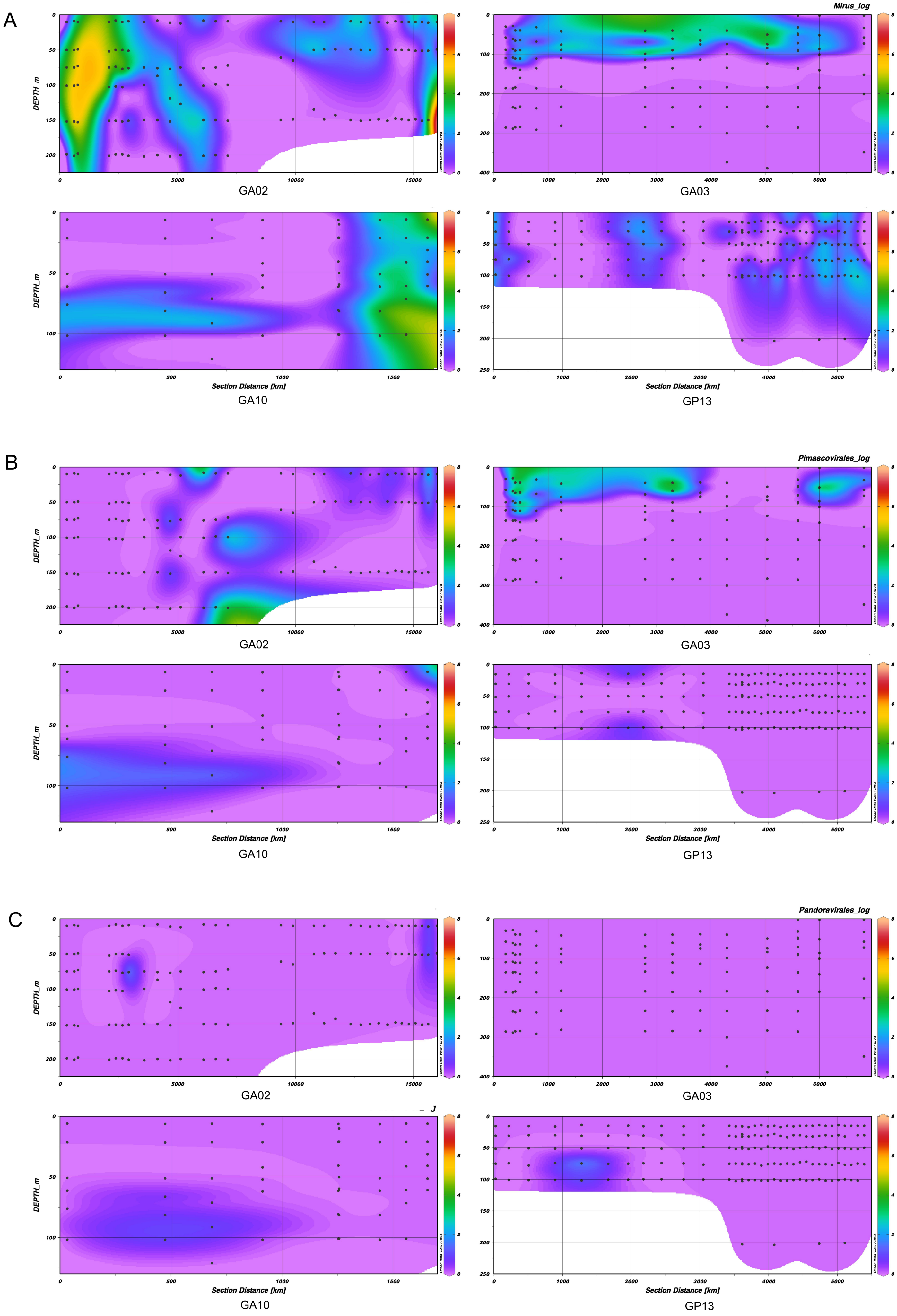
A
Mirus_log
B
Pimascovirales_log
C
Pandoravirales_log
GA02
GA03
GA10
GP13
DEPTH_m
Section Distance [km]
Ocean Data View / DIVA

**Figure S3. Comparison of overall giant virus richness between three depth ranges in the water column across all transects.** Stars showing significant difference between groups (Wilcox test, p-values < 0.05) (* < 0.05, ** < 0.01, *** < 0.001, **** < 0.0001)

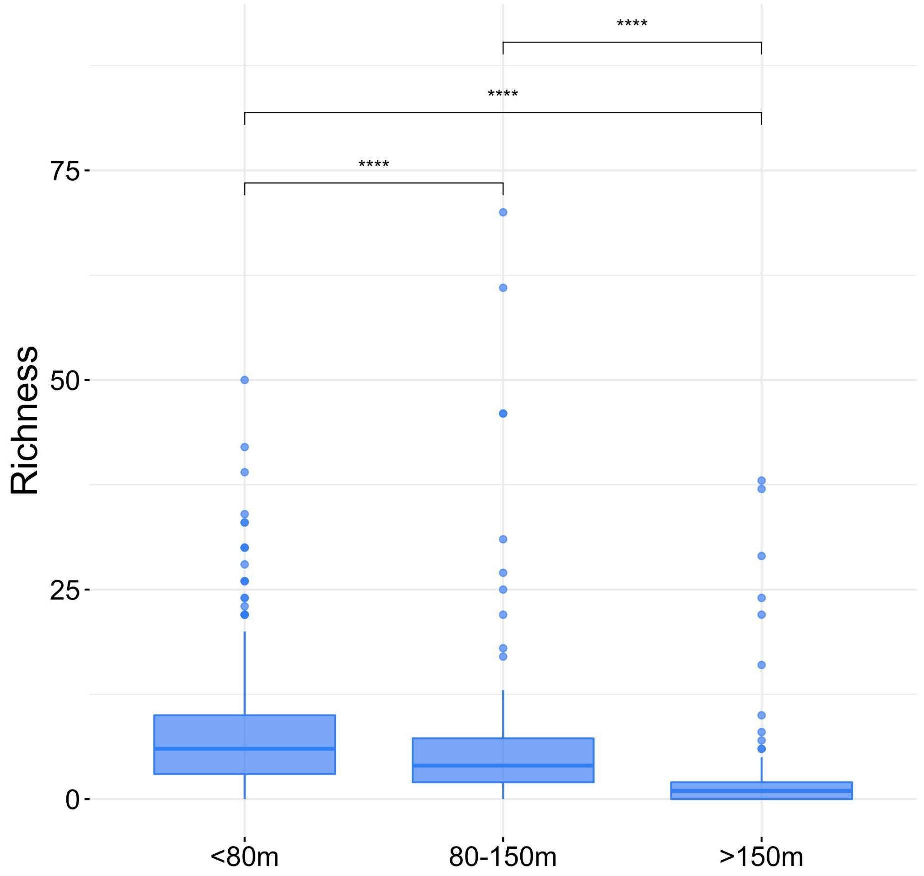

**Figure S4. Relative abundance of giant virus communities along the GA02 transect.** Pie charts indicates the community commposition in each sampling location, with size corresponding to the total abundance of the *Nucleocytoviricota* community.

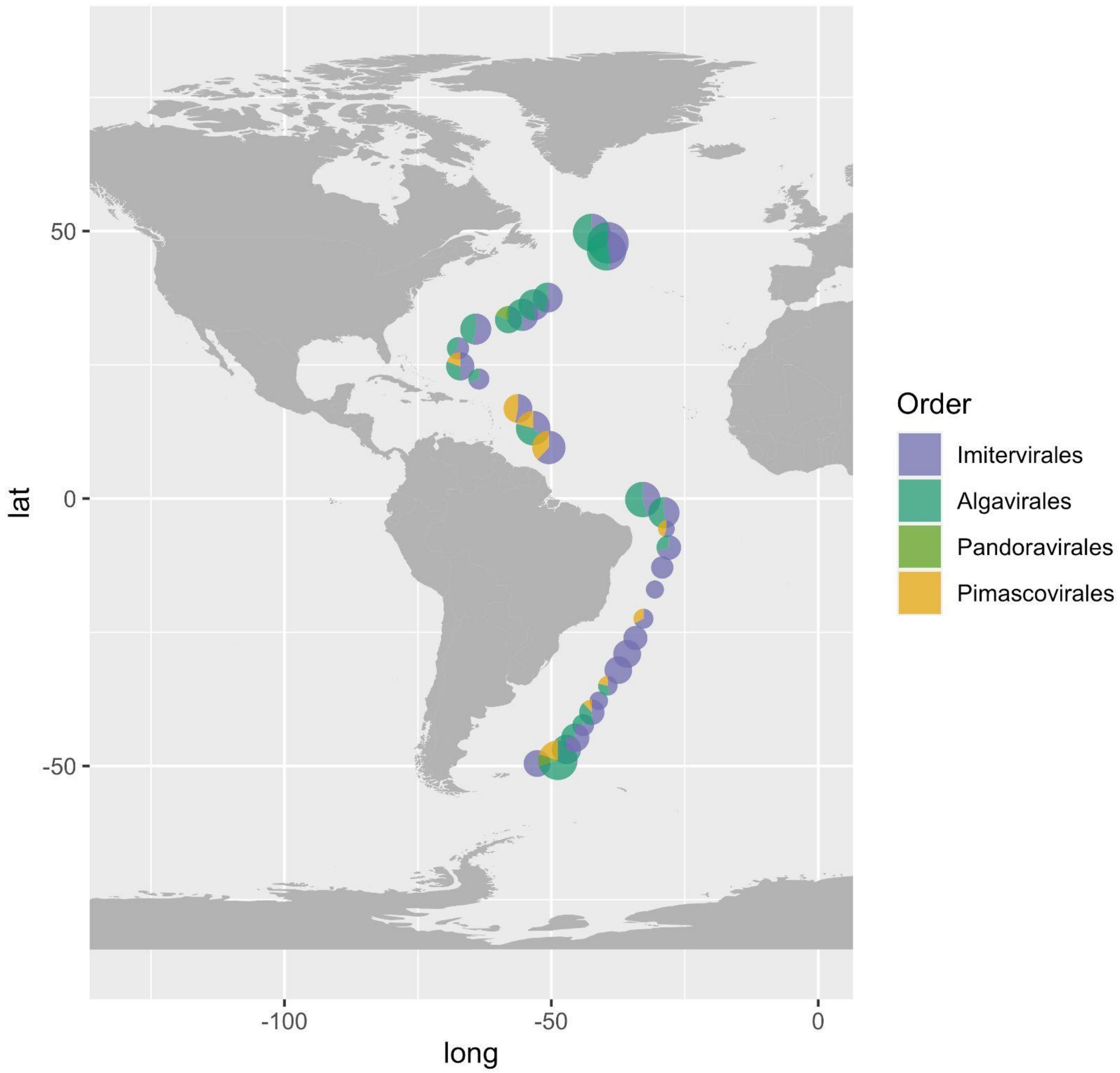

**Figure S5. Latitudinal pattern of giant viral diversity across TARA Oceans samples.** Stars showing significant difference between two latitudinal groups (Wilcox test, p-values < 0.05) (* < 0.05, ** < 0.01, *** < 0.001, **** < 0.0001) Panels left: Shannon's Index; right: Community richness. EQ, Equator.

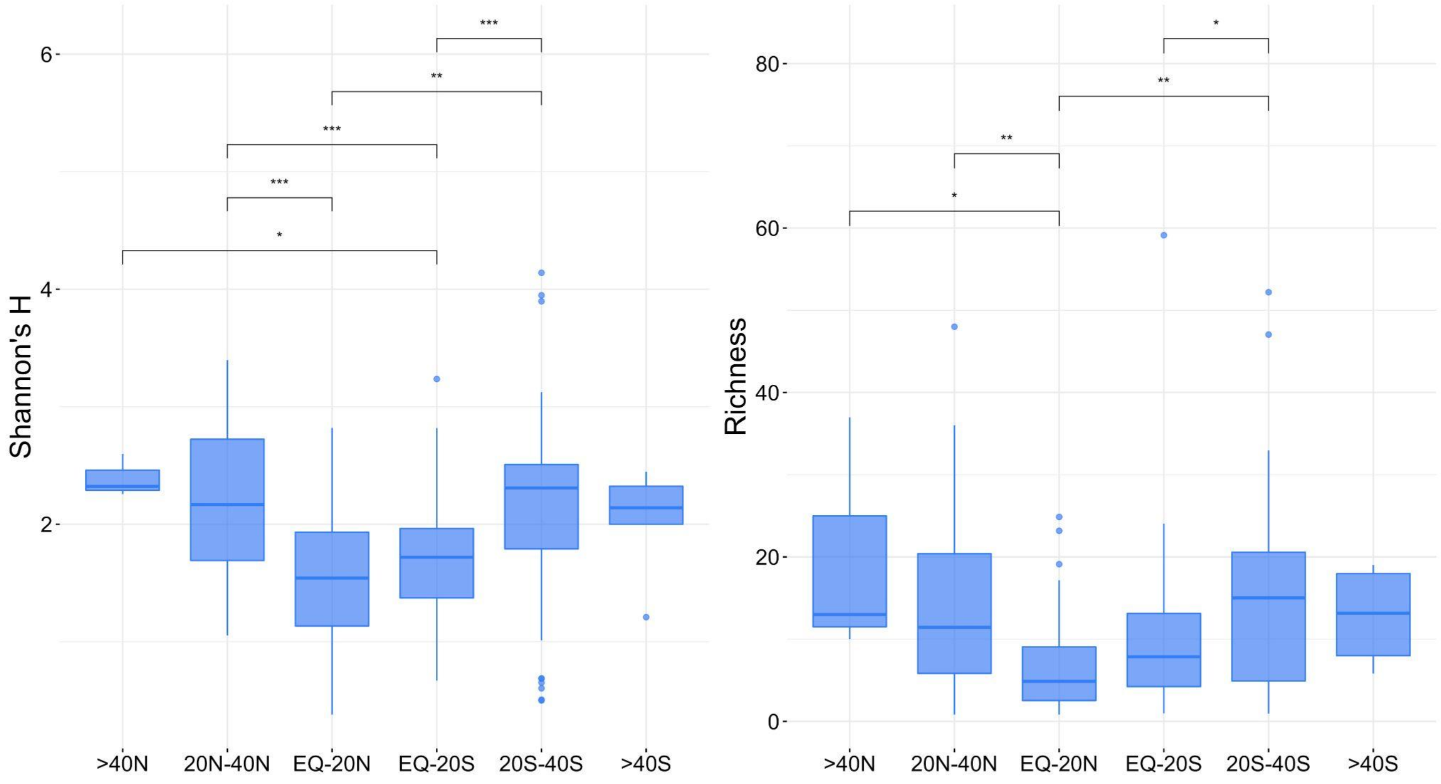

**Figure S6. Unique genomes and genomes shared between the three latitudinal zones.** Horizontal bars (right) indicate the total number of genomes found in each zone; red dots show the average size of all genomes found in each zone. Black dots indicate the presence in one or multiple zones; the corresponding vertical bars indicates the number of genomes with the presence described by the dots.

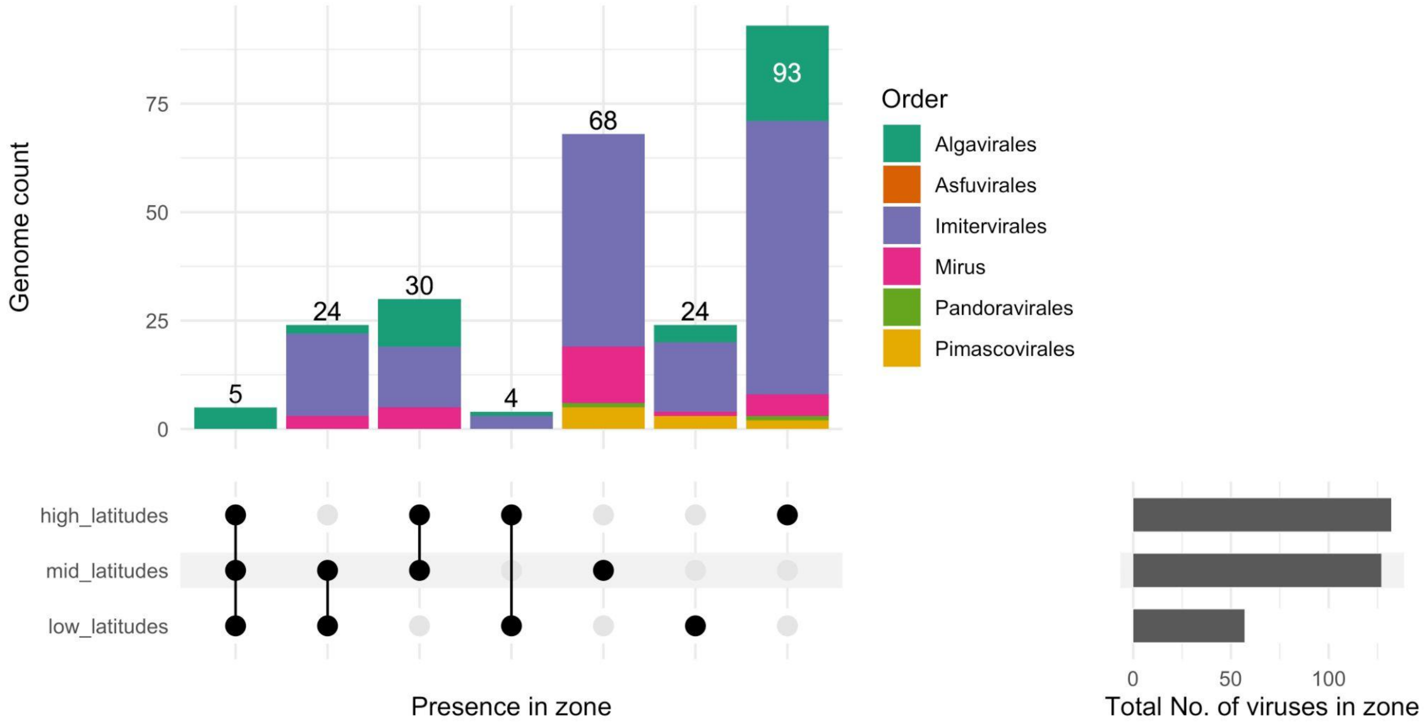

**Figure S7. Geographic distribution of the five *Mesomimiviridae* viruses and one *Prasinoviridae* virus that were widespread in oligotrophic waters.** The size of the bubbles is scaled to the abundance of the virus at a given location. The color of the bubbles shows the taxonomic order of the genome (purple: *Imitervirales*, green: *Algavirales*).

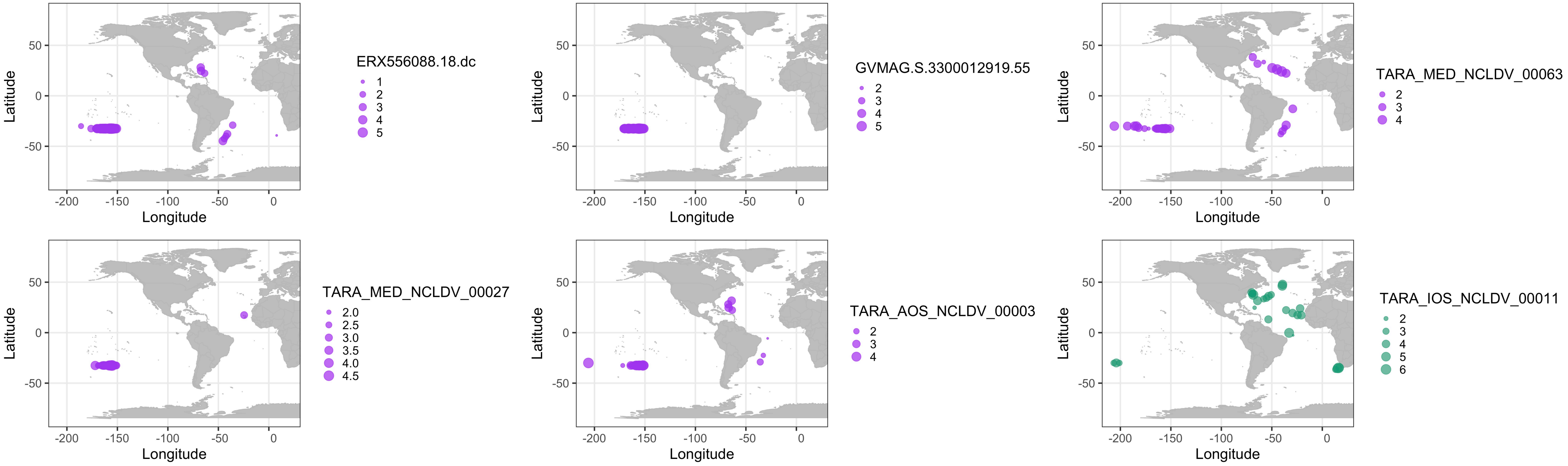
ERX556088.18.dc
1
2
3
4
5
GVMAG.S.3300012919.55
2
3
4
5
TARA_MED_NCLDV_00063
2
3
4
TARA_MED_NCLDV_00027
2.0
2.5
3.0
3.5
4.0
4.5
TARA_AOS_NCLDV_00003
2
3
4
TARA_IOS_NCLDV_00011
2
3
4
5
6
Latitude
50
0
-50
Longitude
-200
-150
-100
-50
0
